# Dorsal striatum and the temporal expectancy of an aversive event in Pavlovian odor fear learning

**DOI:** 10.1101/2021.04.26.441400

**Authors:** Julie Boulanger-Bertolus, Sandrine Parrot, Valérie Doyère, Anne-Marie Mouly

## Abstract

Interval timing, the ability to encode and retrieve the memory of intervals from seconds to minutes, guides fundamental animal behaviors across the phylogenetic tree. In Pavlovian fear conditioning, an initially neutral stimulus (conditioned stimulus, CS) predicts the arrival of an aversive unconditioned stimulus (US, generally a mild foot-shock) at a fixed time interval. Although some studies showed that temporal relations between CS and US events are learned from the outset of conditioning, the question of the memory of time and its underlying neural network in fear conditioning is still poorly understood. The aim of the present study was to investigate the role of the dorsal striatum in timing intervals in odor fear conditioning in male rats. To assess the animal’s interval timing ability in this paradigm, we used the respiratory frequency. This enabled us to detect the emergence of temporal patterns related to the odor-shock time interval from the early stage of learning, confirming that rats are able to encode the odor-shock time interval after few training trials. We carried out reversible inactivation of the dorsal striatum before the acquisition session and before a shift in the learned time interval, and measured the effects of this treatment on the temporal pattern of the respiratory rate. In addition, using intracerebral microdialysis, we monitored extracellular dopamine level in the dorsal striatum throughout odor-shock conditioning and in response to a shift of the odor-shock time interval. Contrary to our initial predictions based on the existing literature on interval timing, we found evidence suggesting that transient inactivation of the dorsal striatum may favor a more precocious buildup of the respiratory frequency’s temporal pattern during the odor-shock interval in a manner that reflected the duration of the interval. Our data further suggest that the conditioning and the learning of a novel time interval were associated with a decrease in dopamine level in the dorsal striatum, but not in the nucleus accumbens. These findings prompt a reassessment of the role of the striatum and striatal dopamine in interval timing, at least when considering Pavlovian aversive conditioning.

## Introduction

Learning time intervals is crucial to survival and goal reaching across the phylogenetic tree. In Pavlovian fear conditioning, an initially neutral stimulus predicts the arrival of an aversive unconditioned stimulus, after a time interval that is encoded, a process pertaining to interval timing (Molet & Miller, 2014; Kirkpatrick & Balsam, 2016). How the brain processes and encodes such information remains poorly understood (Merchant et al., 2013; Tallot & Doyère, 2020). From a neurobiological point of view, there is substantial support for the involvement of the dorsal striatum and its dopaminergic inputs in interval timing (Buhusi & Meck, 2005), both from lesion experiments of the dorsal striatum or intrastriatal infusion of dopaminergic antagonists (De Corte et al., 2019; Meck, 2006), and from lesion or optogenetic manipulation of the substantia nigra pars compacta (SNc), the primary source of dorsostriatal dopamine (Meck, 2006; Soares et al., 2016). Electrophysiological recordings of single neurons in the dorsal striatum of rats also show that striatal neurons firing rate is correlated with the duration of the time interval between signaling cue and reward (Bakhurin et al., 2017; Gouvêa et al., 2015; Matell et al., 2003; Mello et al., 2015), and intrastriatal muscimol infusions produce an impairment in the animals’ ability to discriminate durations (Gouvêa et al., 2015). In the context of fear conditioning, using 2-Deoxyglucose (2-DG) metabolic mapping we previously showed that odor-shock pairing in rats was associated with an increase in 2-DG uptake in the dorsal striatum (Boulanger Bertolus et al., 2014). Furthermore, recent work recording oscillatory neural activity in the dorsal striatum in Pavlovian fear conditioning correlated its maximum power in theta and gamma bands with the time at which the rat expected the aversive stimulus (Dallérac et al., 2017). Timing in fear conditioning is also associated with plasticity in the striatum (Dallérac et al., 2017).

Notably, a majority of studies investigating the neurobiological substrate of interval timing in rodent models rely on operant conditioning. Such protocol relies heavily on the motor response of the subject, which could bias our understanding of the neural substrate of interval timing *per se*. Indeed, the striatum and cortico-striatal inputs are also a neural substrate for motor and procedural learning (Barnes et al., 2005; Koralek et al., 2013; Martiros et al., 2018), and action selection when the task involves temporal discrimination (Howard et al., 2017). To avoid these possibly confounding factors, we used a non-striatum-dependent behavioral measure, the respiratory frequency, to assess the animal’s interval timing ability in a Pavlovian fear conditioning associating an odor to a mild footshock. Indeed, the pattern of respiratory frequency has been shown to be a good index of the animal’s temporal expectation of the shock arrival (Boulanger Bertolus et al., 2014; Dupin et al., 2020; Shionoya et al., 2013). This allowed us to look more closely at the role of dorsal striatum and its dopamine level in the initial acquisition of an interval duration, as well as when a change in this duration is applied. Based on the existing literature, we made two hypotheses 1) inactivating the dorsal striatum should impair timing behavior and its adaptation to a new interval duration and 2) dopamine level in the dorsal striatum should increase when a new interval duration is introduced. We report that, while a transient inactivation of the striatum did alter the expression of the temporal pattern of respiration, and dopamine level in the striatum was indeed modulated during the learning of a new duration, the directions of the effects were the opposite of those hypothesized. These findings prompt a reconsideration of the role of the striatum and striatal dopamine in interval timing, at least when considering Pavlovian aversive conditioning.

## Methods

### Animals

Twenty-five pair-housed and fifteen single-housed male Long Evans rats (Janvier, France) contributed data for experiment 1 and 2 respectively. They weighed 250-300 g at the start of the experimentation, were housed at 23°C under a 12h light–dark cycle, and food and water were available *ad libitum*. All experiments and surgical procedures were conducted in strict accordance with the 2010/63/EU Council Directive Decree and the French National Committee (87/848) for care and use of laboratory animals. The experiments were carried out under the approval of Direction of Veterinary Service (#69000692), and care was taken at all stages to minimize stress and discomfort to the animals.

### Surgery

Details of the procedures can be found in the supplementary methods. Briefly, animals were anesthetized, received subcutaneous local analgesia and were placed in a stereotaxic frame. For experiment 1, rats were implanted bilaterally in the dorsal striatum with stainless steel guide cannulae. For experiment 2, rats were implanted unilaterally (left side) with a guide cannula targeting the dorsal striatum. To assess the specific involvement of the dopaminergic system innervating the dorsal striatum in this task, control rats were implanted in the nucleus accumbens which receives abundant dopaminergic inputs, albeit from another source, the ventral tegmental area (VTA). All animals were allowed two weeks of post-surgical recovery. During this period, the animals’ behavioral state was daily monitored and scored, and if any sign of suffering was detected, the animal was injected with 2-4mg/kg carprofen i.p.

### Experimental apparatus and paradigm

The experimental cages (described in previous studies (Hegoburu et al., 2009, 2011) and detailed in the supplementary methods) consist of a whole-body plethysmograph for Experiment 1 and Plexiglass cylinder for Experiment 2, both customized with tubing in the ceiling connected to a programmable custom olfactometer, and a shock floor. In this setup, rats underwent odor fear conditioning.

For Experiment 1 (Figure 1A), after familiarization to the conditioning cage, the animals received a Conditioning session (ten odor-shock pairings with a 20s odor-shock interval), a Retention test (six odor presentations) and a Shift session (one 20-s odor-shock pairing, followed by nine 30-s odor-shock pairings) at 24h intervals. They were injected with 0.5 µL of either lidocaine (2%, Sigma-Aldrich France, in sterile saline 0.9%, Lidocaine group, n = 13) or saline (Control group, n = 14) just before the Conditioning and Shift sessions. During each session, the animal’s behavior and respiratory rate were continuously recorded for offline analysis.

**Figure 1:**
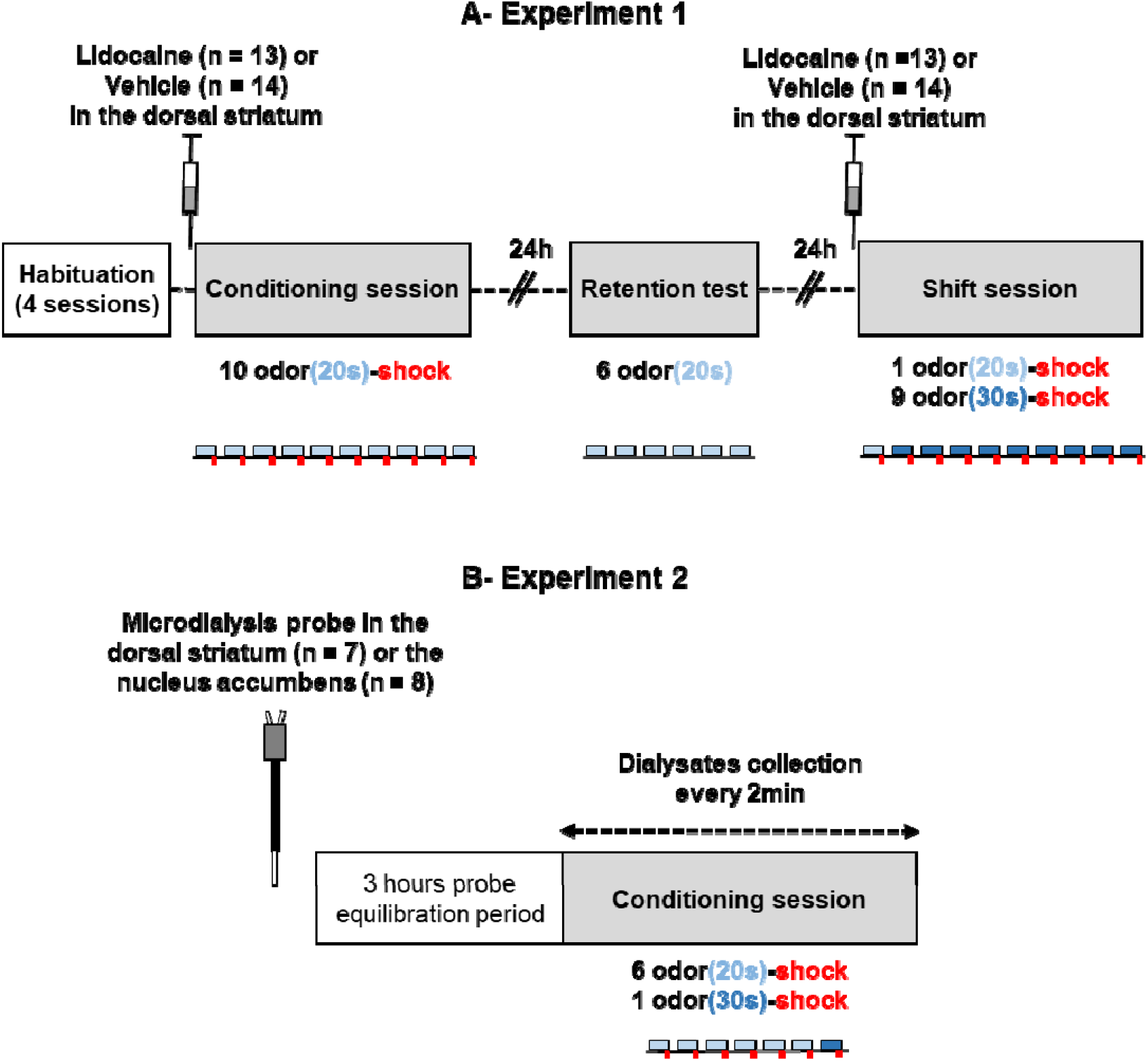
A) Training protocol for Experiment 1. After a period of habituation to the experimental setup for four 20-min sessions, animals were injected with either lidocaine or vehicle in the dorsal striatum, and immediately trained with 10 Odor-Shock pairings using a 20-s odor-shock interval (Conditioning session). Twenty-four hours later, they received 6 presentations of the odor alone to assess their learned fear to the odor (Retention test). Forty-eight hours after conditioning, they were re-injected with the same drug they received previously, and trained in a Shift session where they received 1 odor-shock pairing with a 20-s odor-shock interval followed by 9 odor-shock pairings with a 30-s interval. B) Training protocol for Experiment 2. The microdialysis probe was inserted in the left dorsal striatum or left nucleus accumbens and the animals were placed in the experimental cage for a 3h probe equilibration period after which they were trained with 6 Odor-Shock pairings with a 20-s interval, followed with a single Odor-Shock pairing with a 30-s interval.

For Experiment 2 (Figure 1B), after a 3-hour probe equilibration period, rats received a conditioning session including six 20-s odor-shock pairings, followed by one 30-s odor-shock pairing. Dialysates from the dorsal striatum (n = 7) and the nucleus accumbens (n = 8) were collected every 2 min.

At the end of the experiment, the animals were sacrificed with a lethal dose of pentobarbital for histological verification of the canulae tips localization (Figure S1 and S2 for Experiment 1 and 2 respectively).

### Data acquisition, pre-processing and analysis

In Experiment 1, the respiratory signal and behavior were analyzed as described before (Boulanger Bertolus et al., 2014). We assessed the effects of treatment on the temporal dynamics of the respiratory frequency from the odor onset to shock arrival. For this analysis, the time course of the respiratory frequency, in 1-s time bins, during the 19 seconds of the odor-shock interval was compared using a two-way ANOVA with Group (Lidocaine or Control) as an independent factor, and Time (1 to 19 seconds) as a repeated measure factor. During the Retention test, the freezing rate was analyzed using a two-way ANOVA with Group as an independent factor, and Period (Pre-Odor vs Odor) as a repeated measure factor. For the Shift session, a three-way ANOVA was performed with Group as an independent factor, and Time (1 to 19 seconds) and Interval (20s vs 30s) as repeated measures factors.

In Experiment 2, microdialysis data were acquired as previously described to accurately correlate the neurochemical data with the behavioral events (Parrot et al., 2004; Hegoburu et al., 2009; Hegoburu, Denoroy, et al., 2014), using homemade concentric microdialysis probes (see supplementary methods) continuously infused with artificial cerebrospinal fluid (aCSF). Dialysates were collected in PCR tubes rinsed with an acidic preservative medium, and stored at −30°C until analysis using ultra-high-performance liquid chromatography (see supplementary methods).

Data were expressed as percentage of the baseline obtained by averaging the dopamine concentrations measured in the seven samples collected before the start of the conditioning. Changes in dopamine concentration were then analyzed using a two-way ANOVA with Structure (dorsal striatum or nucleus accumbens) as an independent factor, and Time as a repeated measure factor.

All ANOVA results are reported in the legend of the corresponding figures. For all statistical comparisons performed, post-hoc pairwise comparisons were carried out when allowed by the ANOVA results, the significance level being set at 0.05.

## Results

### Experiment 1: Reversible inactivation of the dorsal striatum during the shift session hastens the adaptation of the respiratory temporal curve to the new interval duration

We previously showed that adult rats submitted to an odor fear conditioning exhibit a typical temporal respiratory frequency pattern (using 1-s time bins) during the odor-shock interval, consisting in a rapid respiratory frequency increase upon odor delivery and a U-shaped decrease just before shock arrival (Boulanger Bertolus et al., 2014; Shionoya et al., 2013). We assessed the effects of dorsal striatum inactivation on this respiratory temporal pattern.

During the Conditioning session, rats with lidocaine-inactivated dorsal striatum exhibited the typical temporal respiratory frequency pattern as early as within the first 3 trials following the first odor-shock presentation (Figure 2A). Within-group comparisons showed that the respiratory rate increased in response to odor arrival in both groups (p < 0.05 from second 4 on), but contrary to the Control group, the Lidocaine group showed a U-shaped decrease preceding shock arrival (significant difference between seconds 8-10 and seconds 15-18). Importantly, the temporal patterns of respiration were similar in both groups by the end of the session, as well as in subsequent odor presentations (Figure S3). Furthermore, during the Retention test, 24h later, both the experimental and control groups showed a significant increase of freezing to the odor compared to the pre-odor baseline (Figure 2B), suggesting a similarly strong odor-shock association in the two groups.

**Figure 2:**
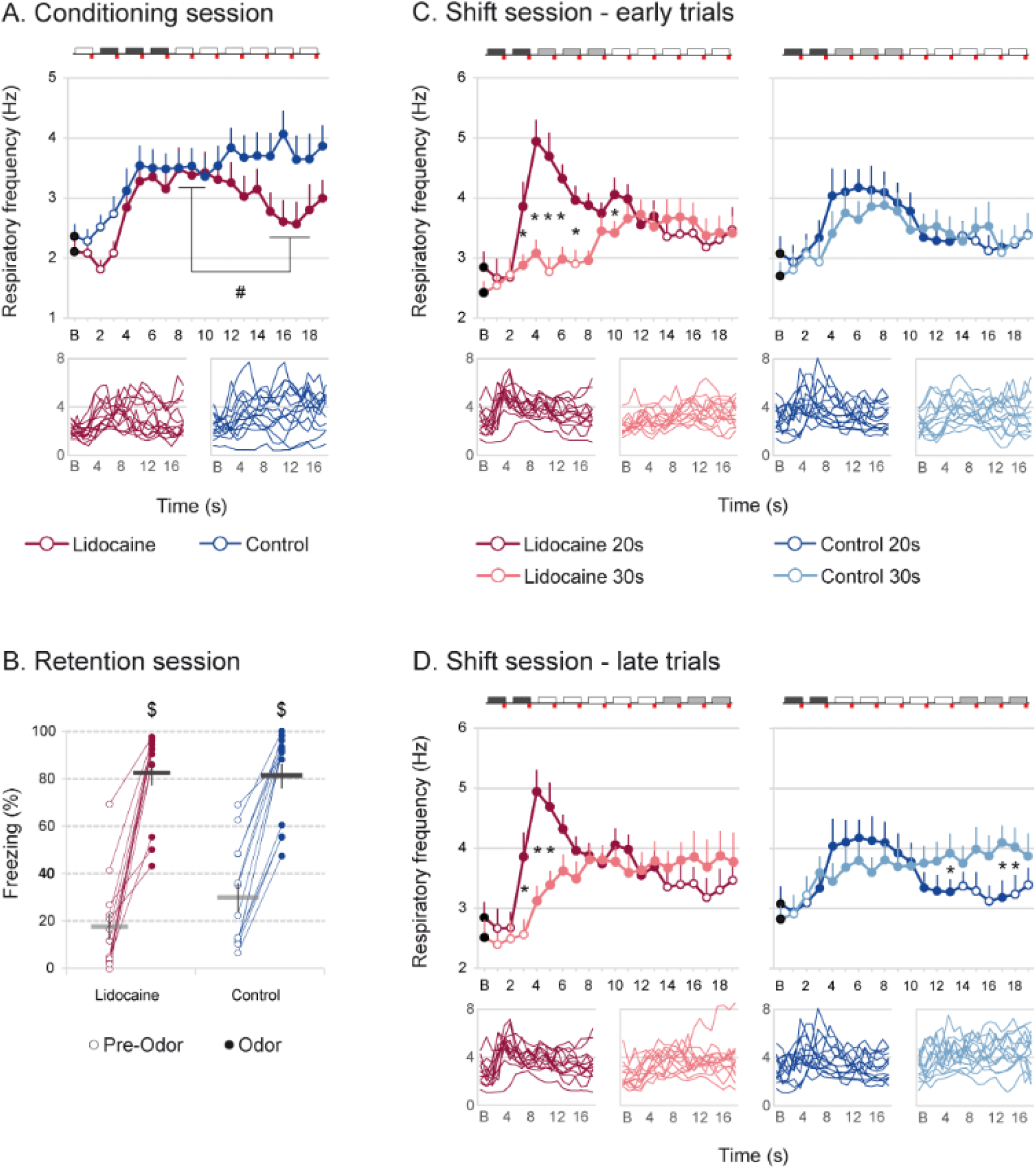
A) Respiratory frequency time course during the 20-s Odor-Shock interval of the Conditioning session, pooled over the 2nd, 3rd and 4th trials (as shown on the schema over the graph), for the Lidocaine (red) and Control (blue) groups. The baseline (B, black dots) corresponds to the average respiratory frequency over the 25s preceding the odor onset. A two-way ANOVA revealed a tendency for a significant Group x Time interaction (F_18,450_ = 1.54, p = 0.07). Filled dots are significantly different from baseline, p<0.05. Inserts represent individual curves. ^#^ Significant difference between designated points, p<0.05. B) Mean percentage of freezing per minute during the 2min period preceding the first odor presentation (Pre-Odor) and during the 6min during which 20-s odor presentations are applied at the beginning of each minute (Odor). The ANOVA confirmed a significant effect of Period (F_1,24_ = 175.4, p < 0.001), but no effect of Group (F < 1) and no significant Period x Group interaction (F_1,24_ = 2.24, p = 0.15). ^$^ Significant difference between Pre-Odor and Odor periods, p<0.05. C) Early shift: Respiratory frequency time course during the 20-s (dark colors; first 2 trials) versus 30-s (light colors; following 3 trials). Odor-Shock interval at the early stage of the Shift session, for the Lidocaine group (Left panel, red) and the Control group (Right panel, blue). A three-way ANOVA confirmed a significant Interval x Time x Group interaction (F_18,450_ = 1.84, p = 0.019). Follow-up analyses showed a significant Interval x Time interaction in the Lidocaine group (F_18,216_ = 8.57, p < 0.001), but not in the Control group (F < 1). (D) Late shift: Respiratory frequency time course during the 20-s (dark colors; first 2 trials) versus 30-s (light colors; last three trials) Odor-Shock intervals at the late stage of the Shift session, for the Lidocaine group (Left panel, in red) and the Control group (Right panel, in blue). The analysis showed an Interval x Time interaction comparing the 20-s trials with the last three trials of the shift (F18,450 = 64.19, p < 0.001), but no Group x Time interaction (F18,450 = 14.57, p = 0.13) nor Group x Interval x Time interaction (F18,450= 12.41, p = 0.54), confirming that both groups shifted their anticipatory response towards the new interval by the end of the Shift session. Filled dots are significantly different from baseline (i.e. black dots, p<0.05). * Significant between intervals difference (p<0.05).

For the Shift session, we compared the mean respiratory frequency temporal pattern obtained during the two first trials (in which the two groups expected the shock at 20s and presented similar respiratory patterns, see Figure S3C) with that obtained during the next three odor-shock pairings during which the animals experienced the shock at 30s. The data showed an early adaptation to the new duration in the Lidocaine group, but not in the Control group (Figure 2C). The scalar rule, a hallmark of interval timing according to which the error in estimating a duration is proportional to the timed duration, predicts a superior superposition of patterns of responses to different time intervals when the time axis is normalized than when it is absolute (Gibbon, 1977). To assess this, the time axis for the 30-s data was multiplicatively rescaled to fit that of the 20-s data (Figure S4) and superposition was indexed by eta-squared (η^2^) as described before (Boulanger Bertolus et al., 2014; Brown et al., 1992; Shionoya et al., 2013). This analysis indicated that the scalar property was respected for the Lidocaine group, but not for the Control group, further confirming that the shift occurred earlier in the Lidocaine group (Figure S4A). Importantly, by the end of the Shift session the temporal patterns of the respiratory response were not different between groups, and the scalar property was respected for both group (Figure S4B).

Together, these data suggest that reversible inactivation of the dorsal striatum hastens an adaptation of the respiratory temporal curve to a newly presented interval duration.

### Experiment 2: Modulation of striatal dopamine level during the acquisition of a new duration

We then monitored dopamine content in dorsal striatum and nucleus accumbens using intracerebral microdialysis with a 2-min sampling rate, during the acquisition session. The session included seven odor-shock pairings, six pairings with a 20-s odor-shock interval and the last pairing with a 30-s odor-shock interval. Such analysis revealed significant differences in dopamine modulation throughout the session, between the dorsal and ventral (nucleus accumbens) striatum (Figure 3A). Structure-specific analyses showed significant modulation of the dopamine level throughout the conditioning in the dorsal striatum, but not in the nucleus accumbens. Post-hoc analysis in the dorsal striatum showed that dopamine level decreased from the beginning of the conditioning, reaching significant differences from baseline from sample 5 (3^rd^ odor-shock presentation) to 10 (except sample 9) after conditioning onset. Dopamine level then slowly re-increased for samples 11 and 12. Interestingly, the last odor-shock pairing for which the interval was shifted to 30s instead of 20s (sample 13) was associated with a significant drop in dopamine level (significant difference with preceding sample, p = 0.046, and following sample, p = 0.016, Figure 3B). More specifically, this drop was observed for 6 out of 7 animals (Figure 3B right panel).

**Figure 3:**
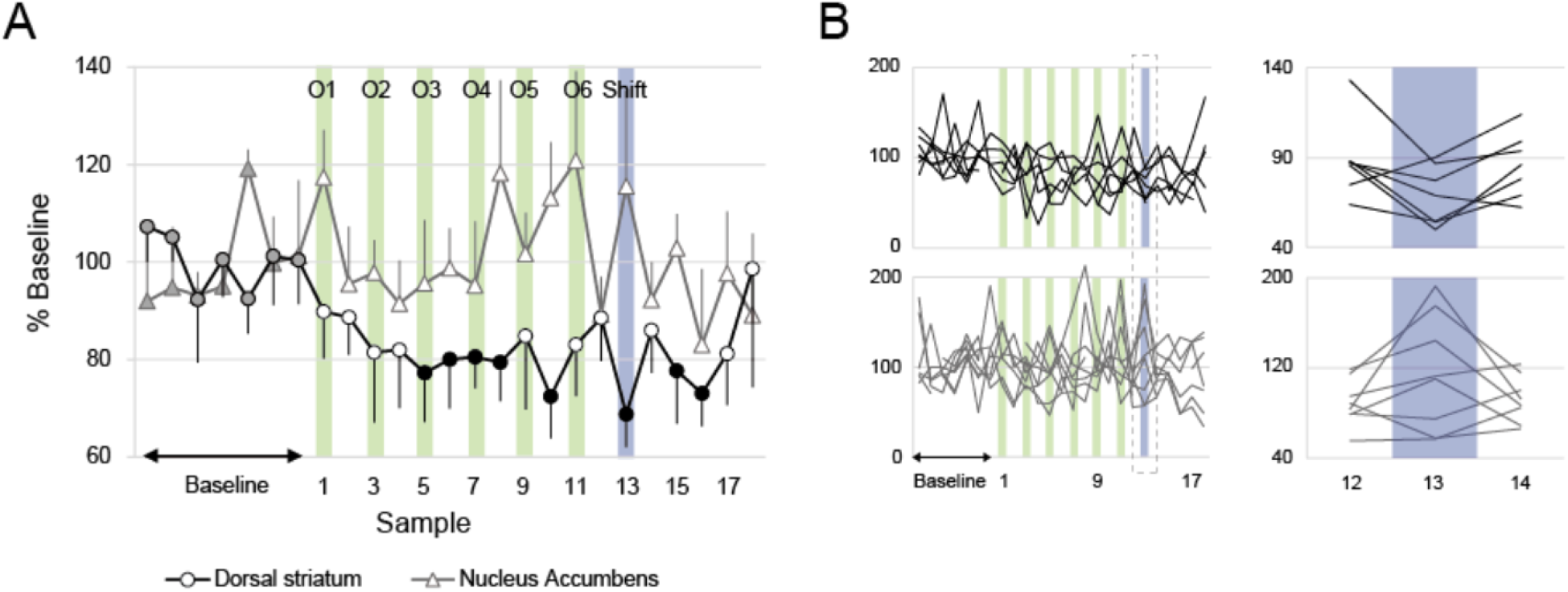
A. Dopamine level in the dorsal striatum (circles, black) and the nucleus accumbens (triangles, grey) during Experiment 2. Dopamine level is measured every two minutes and expressed as a percentage of baseline level measured during the 14min preceding the first Odor-Shock presentation (grey filled symbols). The session included six 20-s Odor-Shock pairings (O1 to O6, green vertical lines) and one 30-s Odor-Shock pairing (Shift, blue vertical line). A two-way ANOVA revealed a significant difference between Structures (F_1,13_ = 9.79, p = 0.008), as well as a significant Structure x Time interaction (F_24,312_ = 1.55, p = 0.050). Further analyses within each structure revealed a significant effect of Time in the dorsal striatum (F_24,144_ = 2.17, p = 0.003), but not in the nucleus accumbens (F_24,168_ = 1.24, p = 0.22). Black filled dots are significantly different from the baseline (p<0.05). B. Individual curves of the dopamine levels in the dorsal striatum (black, top) and the nucleus accumbens (grey, bottom). The right panels show a zoom on the 30-s Odor-Shock pairing (Shift).

## Discussion

In the present study, we investigated how reversible inactivation of the dorsal striatum impacts the learning of novel time intervals using the respiratory frequency to assess male rat’s temporal expectation in an olfactory fear conditioning paradigm. We further investigated the modulations of striatal dopamine extracellular level in that task. Contrary to our hypotheses, we found that reversible inactivation of the dorsal striatum was associated with a hastening of the adaptation of the respiratory frequency temporal pattern during the odor-shock interval. We also showed that the fear conditioning and the learning of a novel interval duration were associated with decreases of dopamine level in the dorsal striatum, while no change in dopamine level was observed in the nucleus accumbens.

In line with the literature (Davis et al., 1989; Dìaz-Mataix et al., 2013; Drew et al., 2005; Dupin et al., 2020; Hegoburu, Parrot, et al., 2014; Laurent-Demir & Jaffard, 2000; Ohyama et al., 2006; Tallot et al., 2020), our results support the assumption that learning a time interval happens early in conditioning, and further suggest that the expression of that learning can be modulated by the dorsal striatum. Indeed, inactivation of the dorsal striatum hastened behavioral adaptation of the animal to a novel duration. Importantly, individual differences observed in response to the shift under lidocaine inactivation (Figure 2C) were not associated with differences in the cannulas’ placement (Figure S1), suggesting they may instead result from interindividual variability in timing abilities (see Dupin et al., 2020 for a discussion of timing abilities and associated brain dynamics). This result suggests that an intact dorsal striatum allows the preservation of the temporal behavioral response pattern to the previously learned time interval at the detriment of the shift of the pattern toward the new duration. This may seem inconsistent with the existing literature. Indeed, electrophysiological recordings in the dorsal striatum during interval timing tasks have consistently shown that the activity of striatal neurons is correlated with the animal’s behavioral temporal response (Bakhurin et al., 2017; Gouvêa et al., 2015; Matell et al., 2003; Mello et al., 2015). Furthermore, lesioning or inactivating the striatum abolishes temporal performance of subjects trained to lever-press for food at a fixed interval (Meck, 2006), or to discriminate between durations (Gouvêa et al., 2015). However, these studies have in common that they rely on operant conditioning tasks that require extensive training of the animal. Importantly, lesions of the dorsal striatum impair the expression of both innate and learned motor sequences (Bailey & Mair, 2006; Cromwell & Berridge, 1996). Consequently, in operant tasks, the specific role of the striatum in the encoding of time cannot be investigated independently from the motor execution of a timed response. Here, using a non-motor task - Pavlovian fear conditioning - and assessing the animals’ behavioral temporal response through respiration, allowed us to overcome this confound, uncovering a previously undescribed role for the striatum. Our results also concur with previous findings indicating that measuring the respiratory response is relevant to understand the brain activity associated with emotions (Dupin et al., 2019; Moberly et al., 2018), including in the context of interval timing (Dupin et al., 2020). As animals were first conditioned to a 20-s interval duration and then shifted to a 30-s duration, one may wonder whether the respiratory patterns observed for these two durations could reflect non-specific changes in performance with trials repetition instead of temporal adaptation to the new duration. However, this is unlikely, as we have shown previously that the respiration curves obtained in animals trained only with a 30-s interval were similar to those of animals shifted to a 30-s interval after training with a 20-s interval as in the present study (Boulanger Bertolus et al., 2014; Dupin et al., 2020).

Importantly, another difference between the present work and previous studies is that most of them used appetitive conditioning to probe the role of striatum in interval timing, while we used an aversive one. Interestingly, Dallérac et al (2017) used a Pavlovian aversive conditioning task, and reported a decreased synaptic plasticity in the striatum when the expected time interval was shifted to a new duration. That study also showed that the amygdala was activated by a shift to a novel interval to time (Dallérac et al., 2017; Dìaz-Mataix et al., 2013), and its inhibition resulted in a faster shift toward the new learned duration. Here, we show that, akin to amygdala’s inhibition, inhibiting the striatum resulted in a speeding of the shift of the behavioral pattern towards the new duration. Together, our data and those of Dallérac et al (2017) suggest that the disengagement of the dorsal striatum, possibly under the control by the amygdala, might facilitate flexible temporal expectancy’s behaviors in Pavlovian aversive conditioning.

Dallérac et al suggested (2017) that this disengagement of the dorsal striatum could be facilitated by dopamine-regulated plasticity (Calabresi et al., 2007; Tritsch & Sabatini, 2012). Using intracerebral microdialysis of dopamine, we were able to show that dopamine level in the dorsal striatum, but not in the nucleus accumbens, decreases when the animal learns the odor-shock association, and when the time interval is changed. A few studies have shown that the dorsal striatum might be involved in fear conditioning (Jeanblanc et al., 2003; Kathirvelu & Colombo, 2013; Matsumoto & Hikosaka, 2009; White & Salinas, 2003). On the other hand, although a few studies have suggested its possible involvement in Pavlovian fear conditioning (Fadok et al., 2010; Wendler et al., 2014), the nucleus accumbens has been mostly implicated in appetitive conditioning (for a review see Schultz, 2016). It might be argued that the observed decreases in dopamine levels in the dorsal striatum were related to non-temporal cognitive processes, such as the mere learning of the odor-shock association or the novelty of the experience. However, changing the interval duration without changing other elements of the association resulted in a further decrease in dopamine level (highlighted in figure 3B), supporting the assumption that dopamine in the dorsal striatum is indeed modulated by the temporal manipulation. Of note, in our study, dopamine level was analyzed every two minutes, which does not allow to ascribe the observed changes to the arrival of the odor or of the shock specifically. Nevertheless, this sampling rate permitted a dynamic assessment of dopamine level that revealed a transient change in response to the shift in duration that had not been described before. These data suggest that the dopamine decrease observed in the dorsal striatum could contribute to support two distinct phenomena: the learning of the association itself (or the development of the behavioral response to that learning), which is associated with a progressive decrease in dorsostriatal dopamine, and the detection of a change in the temporal relationships of the elements of the association, which is associated with a transient dopamine decrease.

## Conclusion

The findings of our study suggest that inactivation of the dorsal striatum hastens the behavioral adaptation of the rat’s respiratory response to the time interval embedded in an aversive conditioning. They further indicate that this behavioral adaptation is likely associated with a decreased release of dopamine in the dorsal striatum. Furthermore, they are in stark contrast to our initial predictions based on the existing literature and suggest a need to rethink the role of the striatum and striatal dopamine in interval timing. This discrepancy further calls attention to the advantage for the field to diversify the tasks and behaviors used to study interval timing in order to discriminate between the brain structures involved in the learning of time intervals and those necessary for the expression of these learned time intervals.

## Acknowledgements

This work was supported by the Centre National de la Recherche Scientifique, by Partner University Funds “Emotion & Time” and by the LABEX CORTEX (ANR-11-LABX-0042) of Université de Lyon, within the program “Investissements d’Avenir” (ANR-11-IDEX-0007) operated by the French National Research Agency. JBB was supported by the Department of Anesthesiology of the University of Michigan when writing this article. The authors warmly thank Ms Ounsa Jelassi-Ben Hellal for her continuous care of animals and Ms Morgane Gallet and Cyrielle Audiane for their technical help.

## Author contributions

JBB, SP and AMM designed the study, did the experiments, carried out the analyses and wrote the manuscript; VD discussed the data, commented and edited the manuscript.

## Supplementary Material

### Supplementary Methods

#### Experiment 1: Inactivation of the dorsal striatum during odor fear conditioning Animals

Data were collected from 27 male Long Evans rats (Janvier, France), weighing 250-300 g at the start of the experimentation (2 animals were used and excluded due to wrong positioning of the injection cannulae). They were housed in pairs at 23°C and maintained under a 12h light– dark cycle (lights on from 7:00 a.m. to 7:00 p.m.). Experiments were performed during the light phase. Food and water were available *ad libitum*. All experiments and surgical procedures were conducted in strict accordance with the 2010/63/EU Council Directive Decree and the French National Committee (87/848) for care and use of laboratory animals. The experiments were carried out under the approval of Direction of Veterinary Service (#69000692), and care was taken at all stages to minimize stress and discomfort to the animals.

#### Surgery

Animals were anesthetized with Equithesin, a mixture of chloral hydrate (127 mg/kg, i.p.) and sodium pentobarbital (30 mg/kg, i.p.), administrated by intraperitoneal injection, and placed in a stereotaxic frame (Stoelting, USA). Before head skin incision, lidocaine (1% solution; Sigma-Aldrich, Saint-Quentin Fallavier, France) was administered subcutaneously for local anesthesia. Rats were implanted bilaterally with stainless steel guide cannulae (23G, Phymep, France) positioned 1.5 mm above the targeted location of the injection needle tips in the dorsal striatum, whose final stereotaxic coordinates from Paxinos and Watson (2014) relative to bregma, were as follows: AP: 0.6 mm; L: 2.8 mm; DV: −4.5 mm from dura. The cannulae were fixed to the skull with dental acrylic cement and anchored with surgical screws. Stylets of the length of the guide cannula were inserted in it to prevent clogging. The animals were allowed two weeks of post-surgical recovery.

#### Experimental apparatus

The apparatus has been described in a previous study (Hegoburu et al., 2011). It consisted of a whole-body customized plethysmograph (diameter 20cm, height 30cm, EmkaTechnologies, France) placed in a sound-attenuating cage. The ceiling of the plethysmograph was equipped with a tower which allowed the introduction of three Tygon tubing connected to a programmable custom olfactometer. Deodorized air flowed constantly through the cage (2L/min), a ventilation pump drawing air out of the bottom of the plethysmograph. When programmed, an odor (McCormick Pure Peppermint; 2 L/min; 1:10 peppermint vapor to air) was introduced in the air stream through the switching of a solenoid valve (Fluid automation systems, CH-1290 Versoix, Switzerland). The bottom of the plethysmograph was equipped with a shock floor connected to a programmable Coulbourn shocker (Bilaney Consultants GmbH, Düsseldorf, Germany). Animal’s behavior was monitored with four video cameras (B/W CMOS PINHOLE camera, Velleman, Belgium) placed at each corner of the sound-attenuating cage.

#### Experimental paradigm

Subjects were submitted to a 7d experimental paradigm consisting of 4d of Habituation to the conditioning cage (20min/day), followed by a Conditioning session, a Retention test and a Shift session (Figure 1A) at 24h intervals. During the Conditioning session, the animals were allowed free exploration for 4min, then received ten odor-shock trials during which the conditioned stimulus (CS), a peppermint odor, was introduced into the cage for 20s, the last second of which overlapped with the delivery of a 0.4 mA foot-shock, the unconditioned stimulus (US), with an intertrial interval of 4 minutes. During the Retention test, after a 2-min odor-free period in the experimental cage (equipped with new visual cues and with a plastic floor to avoid contextual fear expression), the CS was then presented 6 times for 20s with a 1-min intertrial interval. During the Shift session, the animals were re-conditioned by receiving a first odor-shock trial using the previously learned 20s interval, after which nine odor-shock trials were carried out with a new (30s) CS-US interval. During the different steps of the experiment, the animal’s behavior and respiration were continuously monitored and recorded for offline analysis.

#### Pharmacological inactivation of the dorsal striatum

Five minutes prior to the Conditioning and Shift sessions, animals were injected with 0.5 µL of either lidocaine (2%, Sigma-Aldrich France, dissolved in sterile saline 0.9%, injection rate 0.5 µL/min, Lidocaine group, n = 13) or saline (Control group, n = 14). Injection needles (30G) extended 1.5 mm from the tip of the guide cannulae, and were connected via polyethylene tubing to two 10-mL Hamilton microsyringes driven by an automated microinfusion pump (Harvard Apparatus, France). After the injection, the needles were left in position for an additional minute to enable diffusion of the solution into the brain tissue. At the end of the experiment, the animals were sacrificed with a lethal dose of pentobarbital for histological verification of the canulae tips.

#### Data acquisition and pre-processing

The respiratory signal collected from the plethysmograph was amplified and sent to an acquisition card (MC-1608FS, Measurement Computing, USA; Sampling rate = 1000 Hz) for storage and offline analysis. The detection of the respiratory cycles was achieved using an algorithm described in a previous study (Roux et al., 2006). Momentary respiratory frequency was determined as the inverse of the respiratory cycle (inspiration plus expiration) duration, averaged on a second by second basis, and synchronized to the odor arrival and shock delivery using TTL signals. The video signal collected through the four video cameras was acquired with homemade acquisition software using the Matrox Imaging Library and a Matrox acquisition card (Morphis QxT 16VD/M4, Matrox video, UK). Offline, the animal’s freezing behavior was automatically encoded via a LabView homemade software that had been validated by comparison to hand scoring by an experimenter blind to the rat’s group. Data were analyzed using scripts in Python.

#### Data analysis

We assessed the effects of treatment on the temporal dynamics of the respiratory frequency in presence of the CS odor. For this analysis, the time course, in 1-second time bins, of the respiratory frequency during the 19 first seconds of the odor presentation was compared using a two-way ANOVA with the group (Lidocaine vs Control) as an independent factor and the time (seconds 1 to 19) as a repeated measure factor. Post-hoc pairwise comparisons were then carried out when allowed by the ANOVA results. For all statistical comparisons performed, the significance level was set at 0.05.

During the Retention test, the conditioned fear response was assessed by comparing the amount of freezing per minute before and during the odor introduction, using a two-way ANOVA with the group as an independent factor and the period (Pre-Odor vs Odor) as a repeated measure factor. Pairwise comparisons were then carried out when allowed by the ANOVA results.

#### EXPERIMENT 2: Microdialysis during odor fear conditioning Animals

Twenty-one male Long Evans rats (Janvier Labs, France), weighing 250-300 g at the start of the experimentation were used for this experiment (6 animals were excluded for wrong positioning of the probe or technical problem during the session). They were individually housed in the environmental conditions described above.

#### Surgery

The animals were anesthetized with ketamine (70 mg/kg) and xylazine (6 mg/kg) administrated by intraperitoneal injection, and placed in a stereotaxic frame (Stoelting, USA). Before head skin incision, lidocaine (1% solution; Sigma-Aldrich, Saint-Quentin Fallavier, France) was administered subcutaneously for local anesthesia. The rats were implanted unilaterally (left side) with a guide cannula for microdialysis probe (CMA12, Phymep, France) and positioned 3.5 mm above the targeted location of the dialysis membrane tip, whose final stereotaxic coordinates from (Paxinos & Watson, 2014), relative to bregma, were as follows: dorsal striatum AP: 0.6 mm; L: 3.5 mm; DV: −5.7 mm from dura) or nucleus accumbens (AP: 2 mm; L: 1.2 mm; DV: −7.6 mm from dura). The cannula was fixed to the skull with dental acrylic cement and anchored with two surgical screws. Stylets of the length of the guide cannula were inserted in it to prevent clogging. The animals were allowed 2 weeks of postsurgical recovery during which they were regularly handled and habituated to the experimental chamber for 20 min daily during the four days preceding the microdialysis experiment.

#### Microdialysis procedure

Concentric microdialysis probes were constructed in our laboratory from regenerated cellulose dialysis tubing (Spectra/Por hollow fiber; ref #132274, 225 mm O.D., 2 mm as active length, Spectrum Medical Industries) and fused-silica capillary tubing (90 cm and 80 cm long for inlet and outlet, respectively, 40 mm i.d., 105 mm O.D., Polymicro Technology). The body of the probe consisted of a 26-G stainless steel tubing that was glued on a flat probe holder (Harvard) adaptable to the CMA12 cannula-guide. After being flushed, the probes were continuously infused with artificial cerebrospinal fluid (aCSF: 145.0 mmol/L NaCl, 2.70 mmol/L KCl, 1.0 mmol/L MgCl_2_, 1.20mmol/L CaCl_2_, 0.45mmol/L NaH_2_PO_4_, 2.33mmol/L Na_2_HPO_4_, pH 7.4) using a 500-µL Hamilton syringe mounted on an infusion pump (Harvard Model PHD 2000 Infuse). The aCSF was infused at 1 µL/min for the present experiments to avoid ultrafiltration.

Microdialysis on behaving animals requires long tubing (0.9-1.2 m). At a given flow rate, the dead volume of these tubing (i.e. the tubing volume between the dialysis membrane and the outlet of the probe) results in a dead time that must be taken into account to accurately correlate the neurochemical data with the behavioral events (Parrot et al., 2004; Hegoburu et al., 2009; Hegoburu, Denoroy, et al., 2014), estimated to be ∼ 2 min.

#### Experimental apparatus

The apparatus has been described in a previous study (Hegoburu et al., 2009). It consisted of a Plexiglas transparent cylinder (diameter = 21 cm, height = 21.5 cm) with a lateral door placed in a sound-attenuating cage. The ceiling of the cage was perforated with a central aperture allowing the passage of microdialysis tubing and the branching of three Tygon tubing connected to the programmable custom olfactometer described for experiment 1. The bottom of the cage was equipped with the shock floor as described above and connected to an exhaust fan allowing continuous evacuation of the odorant stream from the cage. Animal’s behavior was monitored with four video cameras (B/W CMOS PINHOLE camera, Velleman, Belgium) placed at each corner of the sound-attenuating cage.

#### Experimental paradigm

On the day of the experiment, the microdialysis probe was inserted into the guide-cannula (Dorsal striatum group : n = 7, Nucleus Accumbens group: n = 8) and the animal was introduced in the experimental cage described in experiment 1. After a 3-hour probe equilibration period, the conditioning session was initiated (Figure 1B). Six odor-shock pairings with a 20-s CS-US interval were presented, with an intertrial interval of 4 minutes. An additional pairing was then presented using a 30s CS-US interval. All along the session, dialysates were collected every 2 min in PCR tubes previously rinsed with an acidic preservative medium. The samples were immediately stored at −20°C. Once all the samples were collected, they were transferred into a −30°C freezer until analysis. These procedures permit to limit greatly the degradation of DA from oxidation due to heat, light and the non-acidic aCSF matrix. Of note, even though the paradigm was conducted in a plethysmograph, the microdialysis tubing prevented the sealing of the plethysmograph and thus the collection of the respiratory signal in this experiment.

#### Microdialysis samples analysis

The dialysates were analyzed using ultra-high-performance liquid chromatography (UHPLC). Dopamine, 1-octanesulfonic acid (OSA), triethylamine (TEA), ethylene–diamine–tetra-acetic acid (EDTA) disodium salt, and sodium hydroxide were purchased from Sigma (St. Louis, MO, USA), potassium dihydrogenphosphate and methanol U-HPLC gradient grade from Fisher Scientific (Loughborough, UK). Ultrapure water was produced using a Milli-Q system (Millipore, Bedford, MA, USA). Standard solutions of 1 mmol/L of dopamine were stored at −30°C as aliquots in 0.1 mol/L hydrochloric acid.

The UHPLC system consisted of a Prominence degasser, a LC-30 AD pump and a SIL-30AC autosampler (Shimadzu, Tokyo, Japan). Detection was carried out at 40°C using a Decade II electrochemical detector fitted with a 0.7 mm glass carbon working electrode, a salt-bridge Ag/AgCl reference electrode, and a 25 m spacer (cell volume 80 nL, Antec, Leyden, The Netherlands). Separations were performed at 40°C (in detector oven) using a 100 × 0.32 mm Kappa Hypersil Gold 1.9 m C18 column (Thermo Scientific). The mobile phase, which was adapted from Ferry et al. (2014), consisted of 0.14 mol/L potassium phosphate, 0.1 mmol/L EDTA, 6 mmol/L OSA, 0.01% TEA (v/v), pH adjusted to 5 with 10 mmol/L sodium hydroxide, 6% methanol, filtered through a 0.22 m cellulose acetate membrane before elution at 8.5 µL/ min. Analytes were detected at an oxidation potential of 700 mV versus the reference electrode. Chromatograms were acquired at a rate of 10 Hz using Lab Solutions software. The acquisition time was 22 min. The injection volume was 1 µL. The sample analysis started the same day of the microdialysis collection to prevent dopamine degradation as much as possible, as 25 samples were collected per animal. In some rare cases of chromatographic matters, the analysis was postponed, but not later than 2 days after collection. On the day of analysis, the samples were placed in the autosampler and kept at +4°C before injection. Concentrations of dopamine were calculated using a day calibration curve. Data were expressed as percentage (mean ± SEM) of the baseline obtained by averaging the dopamine concentrations measured in the seven samples collected before the start of the conditioning session. Changes in dopamine concentration were then analyzed using a two-way ANOVA with the structure (dorsal striatum or nucleus accumbens) as an independent factor and the time as a repeated measure factor. Post-hoc pairwise comparisons were then carried out when allowed by the ANOVA results.

### Supplementary Figures

**Figure S1:**
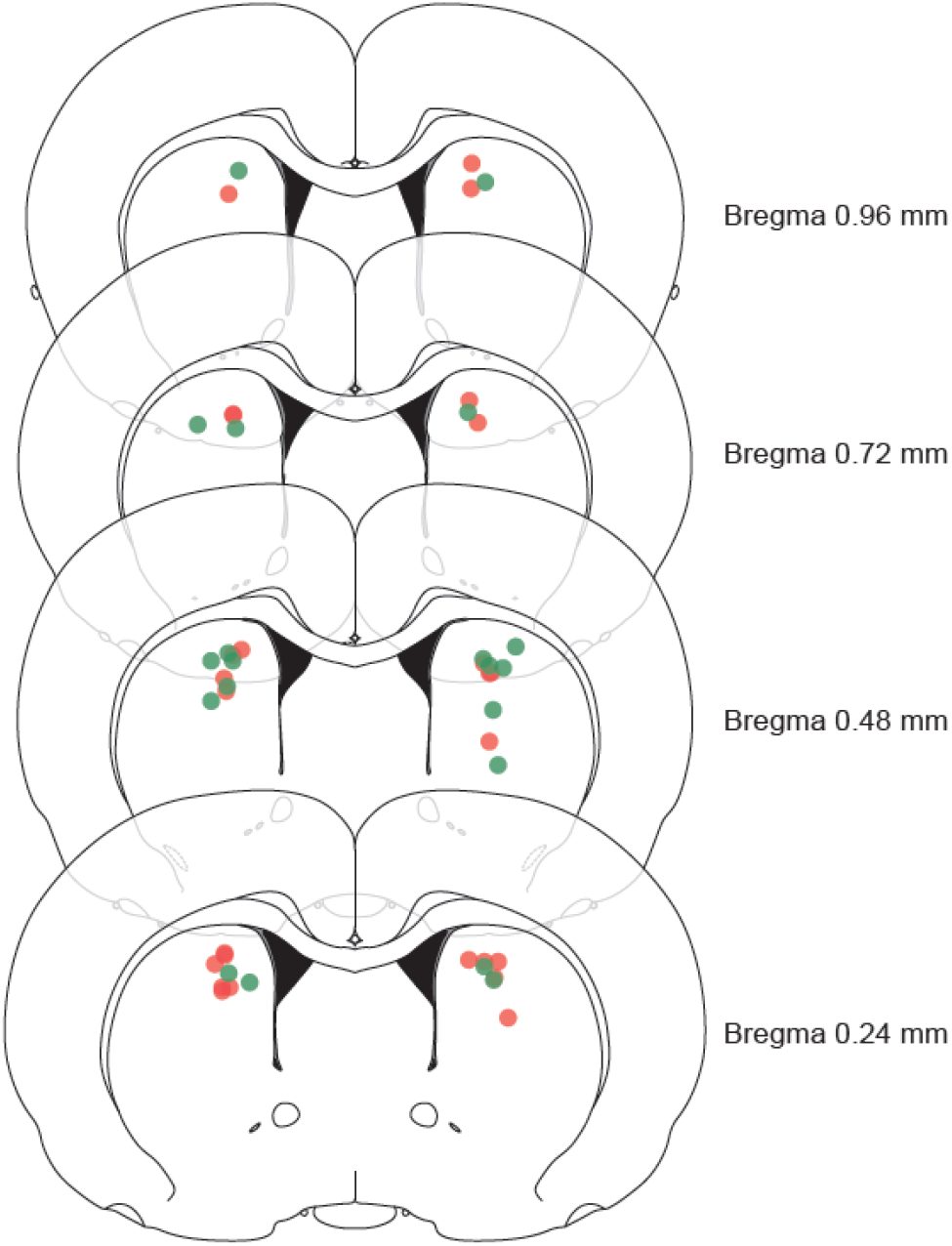
Localization of the tip of the injectors in Experiment 1 for rats in the Lidocaine group (red) and the Control group (green).

**Figure S2:**
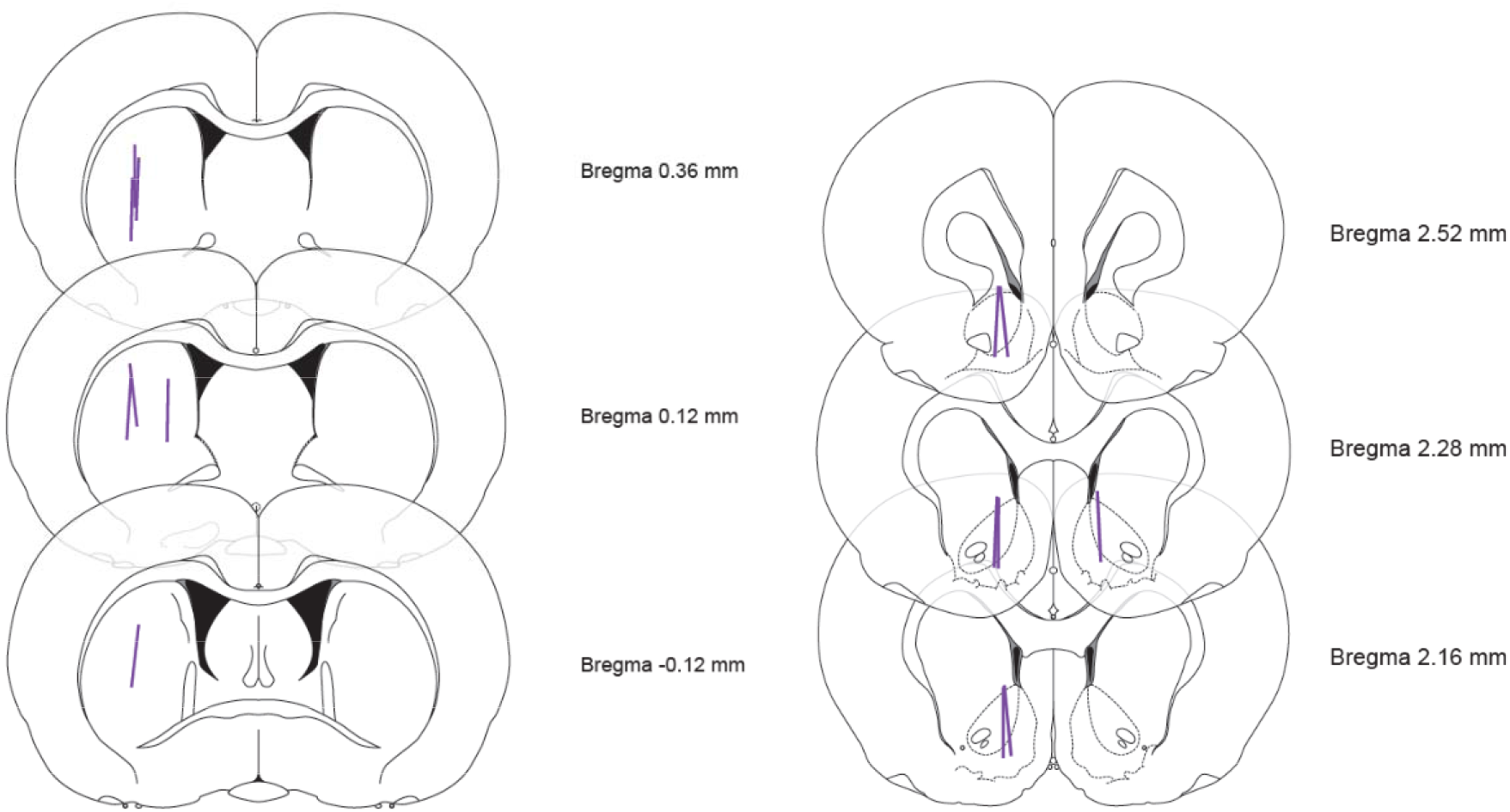
Localization of the microdialysis membranes in Experiments 2 for the rats implanted in the dorsal striatum (left panel) and the nucleus accumbens (right panel).

**Figure S3:**
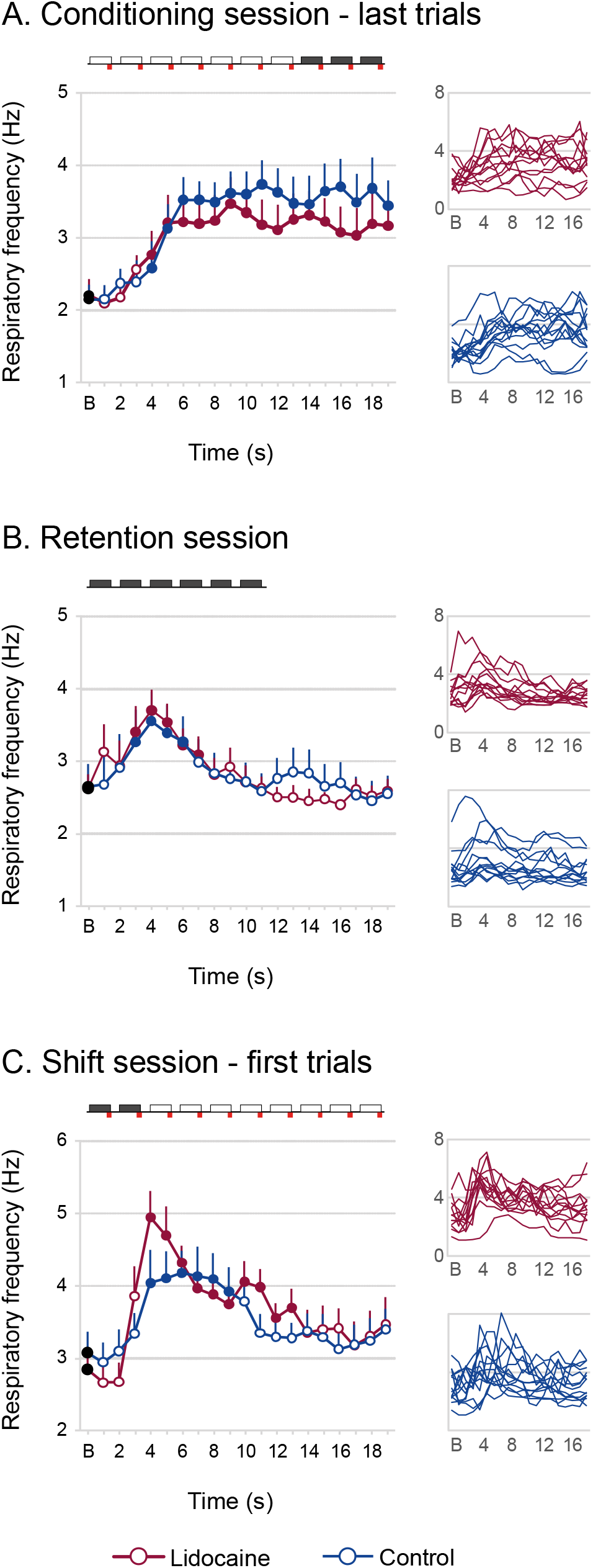
Time course of the respiratory frequency during (A) the last 3 trials of the conditioning session, (B) the 6 trials of the retention test, (C) the first 2 trials of the shift session (as shown on the schema over the graphs), for the Lidocaine (red) and Control groups (blue). ANOVAs with independent factor Group and repeated factor Time show no Group x Time interaction for the last three trials of the Conditioning session: F <1, nor the retention test: F_18,432_ = 1.08, p = 0.37; or for the 2 first trials of the Shift session: F_18,450_ = 1.16, p = 0.29. Filled dots are significantly different from baseline (p<0.05), inserts are the respiratory frequency of individual rats.

**Figure S4:**
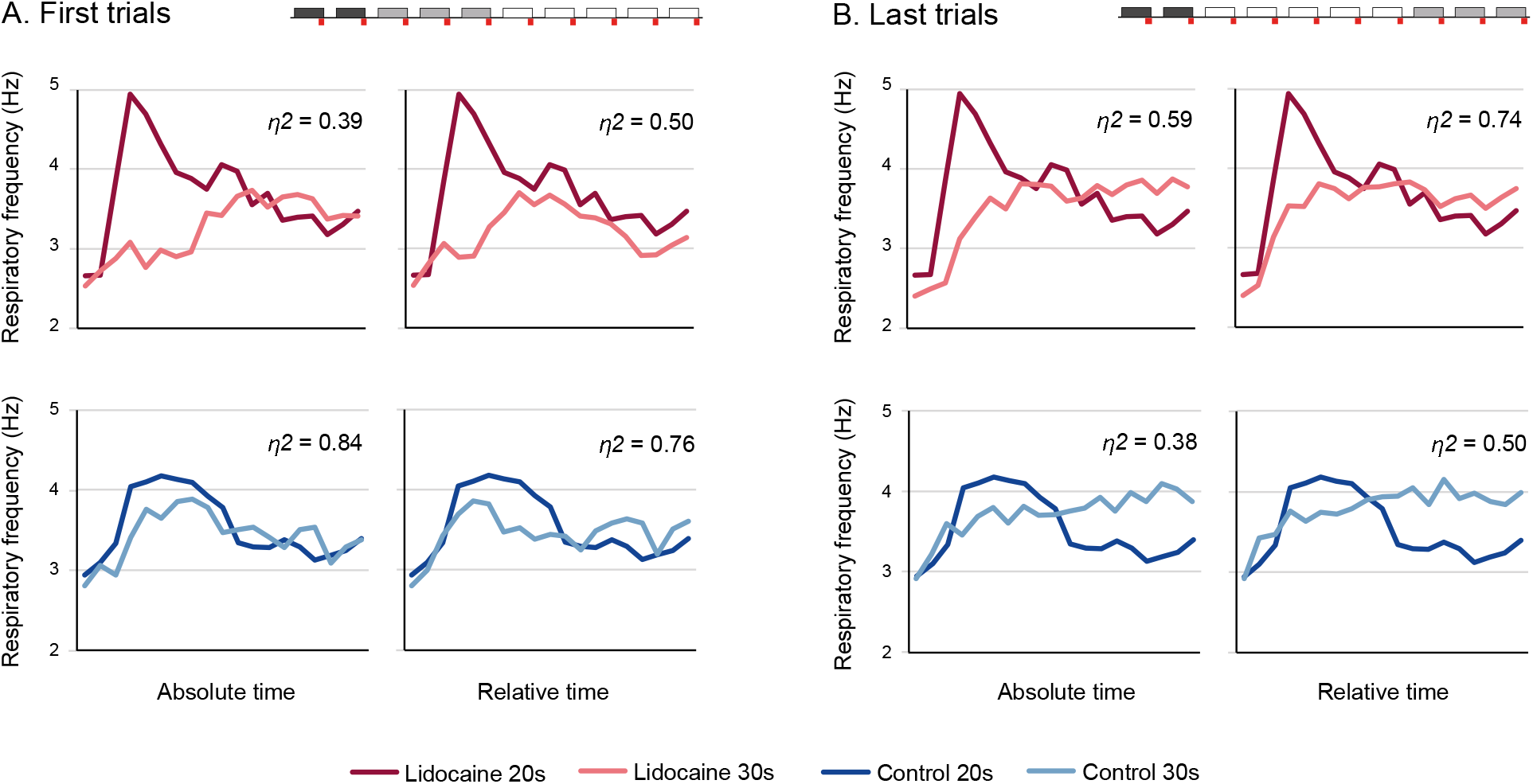
Testing the scalar property of the respiratory frequency curve during the 20-s (in dark colors; first 2 trials) versus 30-s (in light colors) Odor-Shock interval at (A) the early stage of the Shift session of Experiment 1 (first 3 trials after the rats have been exposed to the 30s interval), and (B) the last trials of the shift, for the Lidocaine group (upper panel, red) and the Control group (lower panel, blue). In order to assess scalar timing quantitatively, in each group, the time axis 30-s Odor-shock interval curves was multiplicatively rescaled to superpose with the 20-s interval (relative time). Superposition between the two curves was indexed by eta-square (η2), compared with the superposition index of non-rescaled curves (absolute time), and indicated in the upper right corner of each graph.

